# Differential modulation of thermal preference after sensitization by optogenetic or pharmacological activation of heat-sensitive nociceptors

**DOI:** 10.1101/2020.09.25.312108

**Authors:** Jerry Li, Maham Zain, Robert P. Bonin

**Affiliations:** Departments of Human Biology: Neuroscience and Immunology, University of Toronto, ON, Canada; Leslie Dan Faculty of Pharmacy, University of Toronto, ON M5S 3M2, Canada; University of Toronto Centre for the Study of Pain, University of Toronto, ON M5S 3M2, Canada

**Keywords:** Behavioral analysis, thermal preference, optogenetics, TrpV1

## Abstract

Common approaches to studying chronic pain in pre-clinical animal models paradoxically involve measuring reflexive withdrawal responses that are more indicative of acute nociceptive pain. These methods typically do not capture the ongoing nature of chronic pain nor report on behavioral changes associated with pain. In addition, data collection and analysis protocols are often labour-intensive and require direct investigator interactions, potentially introducing bias. In this study, we develop and characterize a low-cost, easily assembled behavioral assay that yields self-reported temperature preference from mice which is sensitive to peripheral sensitization protocols. This system uses a partially automated and freely available analysis pipeline to streamline the data collection process and enable objective analysis. We found that after intraplantar administration of the TrpV1 agonist, capsaicin, mice preferred to stay in cooler temperatures than control injected mice. We further observed that gabapentin, a non-opioid analgesic commonly prescribed to treat chronic pain, reversed this aversion to higher temperatures. We further observed that optogenetic activation of the central terminals of TrpV1+ primary afferents via *in vivo* spinal light delivery did not induce a similar change in thermal preference, indicating a role for peripheral nociceptor activity in the modulation of temperature preference. We conclude that this easily produced and robust sensory assay provides an alternative approach to investigate the contribution of central and peripheral mechanisms to pathological sensory processing that does not rely on reflexive responses evoked by noxious stimuli.

## Introduction

Chronic pain is pain that persists for more than three months without clear protective benefits and can be a disease by itself (primary chronic pain) or arise from another underlying health condition (secondary chronic pain). These conditions may be precipitated by a series of events or a combination of various risk factors that affect multiple dimensions of an individual’s daily life, including their emotional wellbeing, ability to perform daily tasks, and functioning in the workplace.^1–3^ On a cellular level, pathological pain can arise from sensitization of the peripheral and central nervous systems’ nociceptors.^1, 4^ Injuries and persistent inflammation in the periphery can then lead to central sensitization, characterized by an increase in responsiveness of nociceptive neurons to noxious stimuli.

The mechanistic and behavioral examination of acute nociceptive and pathological pain in preclinical animal models is critical to the understanding of chronic pain pathology and development of novel pain therapeutics.^5^ For many years now, little translational success has placed current paradigms of pain into question.^5, 6^ In preclinical studies of mice, researchers cannot directly assess pain perception in the animals and are limited to quantifying indirect behavioral or physiological measures of pain. The most widely used assays for nociception measure acute pain responses by exposing physically restrained mice to noxious stimuli to elicit reflexive, nocifensive behaviors, such as limb withdrawal, grooming, or paw licking. Typical stimulus-evoked pain assays include the Hargreaves test, von Frey filaments, and cold or hot plate tests.^7.8^ However, these methods are often labour-intensive and require extensive experimenter intervention, thus lending them to increased subjectivity.^9^ Moreover, reflexive nocifensive responses can be purely spinal cord-mediated and can still be seen in anesthetized animals^7, 10, 11^, and may not accurately reflect chronic pain conditions that involve complex supraspinal sensory integration. These concerns have led to increased usage of unconditioned or non-evoked behavioral responses with the aim of capturing endogenous indicators of spontaneous or ongoing pain.^11^ Such measures include operant assays, the Mouse Grimace Scale^12, 13^, and automated behavioral tracking^7^, weight-bearing or gait analysis, nesting or burrowing behavior, ultrasonic vocalizations, and free-choice temperature preference.^14, 15^ Along with increased adoption of assays for spontaneous pain, there has also been an increase in the automation of data collection. Automated analyses allow for reduced subjectivity by minimizing animal handling and removing the need for physical restraints, thereby reducing stress on the animals.

The automated assessment of thermal preference provides a non-invasive approach to investigate peripheral and central processes governing central and peripheral mechanisms of nociception and sensitization. Thermal sensitization can arise following activation of nociceptors expressing the transient receptor potential cation channel subfamily V member 1 (TrpV1) protein that is activated by noxious heat (>43°C), chemicals, and acidity. Thermal sensitization is accompanied by a change in the thermal preference of animals towards cooler temperatures.^17–19^ Capsaicin, a TrpV1 agonist, has been shown to produce significant thermal allodynia and hyperalgesia in both humans and animals via activation of TrpV1.^20^ TrpV1 expression in the adult mouse is mostly restricted to peptidergic C-fibers, a primary afferent subtype that is known to be critical for the relay of nociception.^21^ In addition to capsaicin, recent strides in the development of transgenic animals and optogenetics have allowed precise optical control of TrpV1-expressing afferents. Specifically, in mice expressing the light-activated excitatory ion channel channelrhodopsin (ChR2) in TrpV1^+^ primary afferents, the delivery of blue light to the periphery induces nocifensive behaviors and withdrawal from the light stimulus.^22, 23^ The majority of optogenetic pain research has used transdermal light to activate the primary afferents; while this is a non-invasive strategy, it restricts the delivery of light to the periphery and can complicate experiments involving concurrent optogenetic and thermal or mechanical stimulation.^22, 24^ The development of novel surgical techniques enables non-restrictive optogenetic control of defined afferents and the study of how defined nociceptor populations contribute to nociceptive processing and pain-associated behavior.^25–27^ The multiple approaches available to induce and investigate thermal sensitization in animals allows complementary approaches to be used in the investigation of thermal sensitization.

Given the need for improved approaches to non-invasively study nociceptive processing and spontaneous pain behaviors, we created an automated assay for the assessment of thermal preference in CD-1 and C57BL/6 mice and provided validation of the assay using pharmacological and optogenetic modulation of activity of the transient receptor potential vanilloid 1 (TrpV1) expressing nociceptors. We hypothesized that activation of these nociceptors would result in aversion to higher temperatures and shift temperature preference to cooler temperatures. Our findings provide a basis for further development of automated pain assays and highlight the ability for automation to provide objective, quantitative, easily characterizable, and highly robust data without invasive handling of experimental subjects.

## Materials and Methods

### Thermal preference arena development and calibration

A linear temperature gradient was created across a thin, 40.6 × 10.2 cm plate of aluminum metal (**Fig. 1A**). Three 40 mm × 40 mm 50 W Peltier thermoelectric devices (TEC1-1270640; Hubei I.T., Shanghai) were used to create the gradient. Two Peltier devices were mounted beside each other on one end of the aluminum sheet to provide cooling, while another was mounted on the opposite end in the opposite configuration to heat the aluminum sheet and create the gradient. The Peltier devices were powered by a 480 W computer power supply, with the 5 V/40 A line powering the single warming device, and separate 12 V/35 A power lines used for each cooling device. The Peltier devices providing cooling were mounted on 6.5 cm × 6.5 cm × 6.5 cm aluminum heat sinks. Heat was dissipated from the heat sinks via two 6 cm cooling fans powered by the power supply. The level of noise produced by the power supply and fans cooling the Peltier devices were measured for 5 minutes at stable temperatures using the iPhone app, Decibel X, version 7.0.0. The cold, middle, and hot ends averaged noise levels of 70.5, 68.3, and 67.9 dB, respectively. Four walls of plexiglass were then used to create a 36.0 × 7.4 cm arena atop the aluminum in which mice could freely explore. Black Bristol board surrounded the arena to prevent external distractions. Lastly, a Sony HDR-CX405 video camera was set up directly above the arena to record 30-minute trials. Before each trial, the arena’s temperature gradient was calibrated by turning on the Peltier devices for 15 to 20 minutes and temperature measurements were taken with an infrared thermometer at several points along the aluminum sheet.

**Figure 1.**
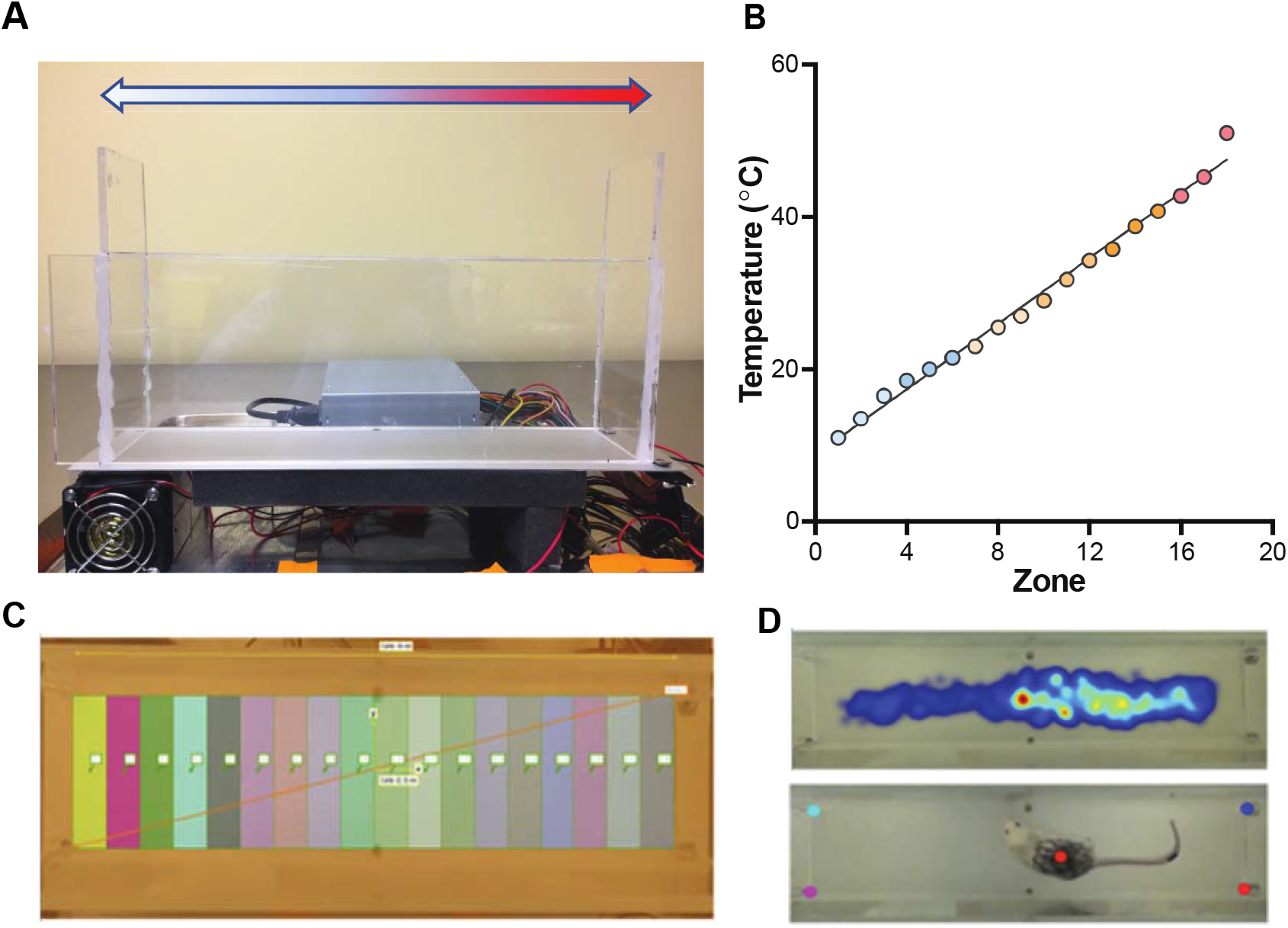
Automated assay setup and tracking parameters. **(A)** A profile of the arena in which mice were placed, tracked with a camera directly above. Peltier devices, powered by a 480 W computer power supply, on either ends of an aluminum plate maintained a linear temperature gradient from 11.0 ± 0.6 (blue) to 51.0 ± 0.7 °C (red). **(B)** Linear temperature gradient (*r*^2^ = 0.89), with each point, or zone, corresponding to measurements taken with an infrared thermometer every 2.0 cm of the arena. **(C)** Eighteen evenly spaced, arbitrarily coloured, 2.0 cm divisions across the arena that correspond to different temperature zones tracked by EthoVision. **(D) Top:** EthoVision’s heatmap analysis of mouse movement throughout the arena in one trial. Mouse position was determined by the location of their centre, as shown in the bottom image. Warmer temperatures correspond to longer time spent in those regions. **Bottom:** DeepLabCut tracking the mouse’s centre in the same trial. Four points on each corner of the arena were used to calculate the position of the mouse.

### Animals and housing

All experiments were conducted in accordance with the Canadian Council on Animal Care and the University of Toronto’s Animal Care and Use guidelines. Adult (>8 weeks old), male CD-1 mice were used for the capsaicin-induced sensitization experiments. 7-week old male CD-1 mice were supplied by Charles River Laboratories and housed in identical enriched home cages in groups of 3-4 upon arrival. They were given a week to habituate to the housing facility, with a 23.0 ± 0.5 °C ambient temperature, 53 ± 13% relative humidity, and 14 h light: 10 h dark cycle. Food and water were given *ad libitum* in their home cages throughout experiments. Individual mice were marked on their tails for identification and their backs for video tracking. Initial marking was done approximately 24 hours before trials began and any re-marking, if necessary, at least 30 minutes before each trial. The mice habituated to the arena itself for 30 minutes before each trial. Experiments were conducted on mice 8-16 weeks of age during the light cycle.

Adult male (>12 weeks old) TrpV1-ChR2 mice were used for the optogenetic sensitization experiments and adult male (>12 weeks old) C57BL/6 mice were used to study the effects of the surgical strategy on mouse wellbeing. All mice were kept in groups of 1 to 4 mice per cage prior to ferrule implantation and singly housed after the implantation surgery with food and water provided *ad libitum.* Mice expressing ChR2 in TrpV1+ nociceptive afferents (TrpV1-ChR2) were generated by crossing mice expressing Cre-recombinase in TrpV1+ neurons with mice expressing a loxP-flanked STOP cassette upstream of a ChR2-EYFP fusion gene at the Rosa 26 locus (Rosa-CAG-LSL-ChR2(H134R)-EYFP-WPRE; stock number 012569, The Jackson Laboratory, Bar Harbor, ME).

### Capsaicin and gabapentin treatments

Two separate randomized crossover studies were conducted. For the first set of trials, 30 minutes after an ipsilateral hind paw injection of either saline (5 μL 0.9%, control) or capsaicin (5 μL, 0.5% w/v; 1:1:8 as 95% ethanol:Tween 20:0.9% saline), each mouse was recorded in the arena for 30 minutes (**Fig. 2A**). For the second set of trials, each mouse first had an intraperitoneal injection of either saline (control) or gabapentin (10 mL/kg), and then after 30 minutes, an ipsilateral hind paw injection of capsaicin of the same concentration. After another 30 minutes, they were recorded in the arena for 30 minutes. All capsaicin injections were conducted after placing the mice under light isoflurane anesthesia, with toe-pinch reflex present. By the end of each set of trials, each mouse served as their own control and had injections of both control and treatment, with a 48-hour washout between each treatment. For each mouse, after the initial control or treatment injection, subsequent trial injections were done in the contralateral hind paw. All capsaicin and gabapentin injections were done using a 50 μL Hamilton syringe and 1 mL syringe, respectively, with 30-gauge needles.

**Figure 2.**
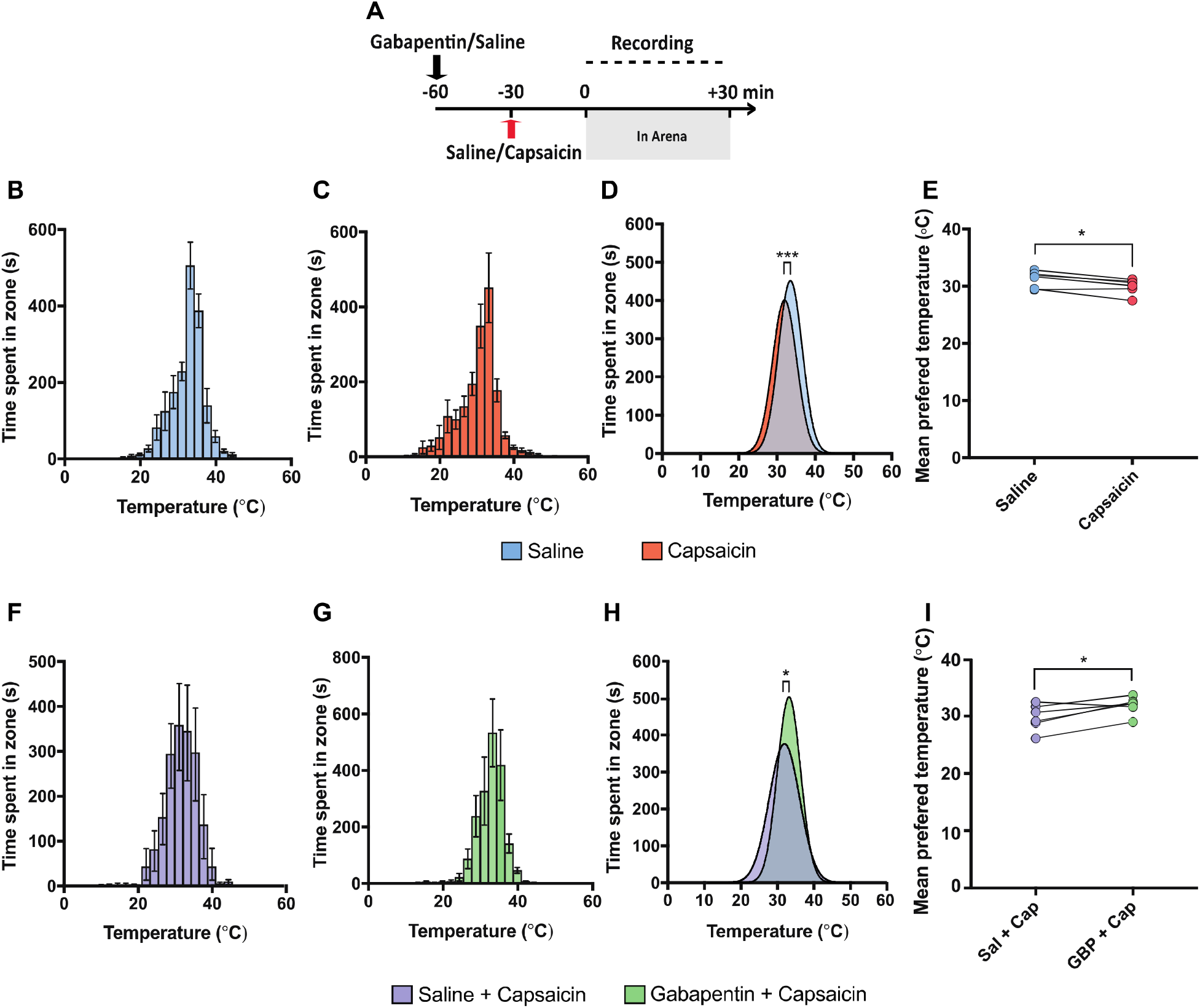
Gabapentin prevented capsaicin-induced change in thermal preference. **(A)** Experimental timeline for the capsaicin-induced sensitization trials. **(B, C, F, G)** Total time spent in each temperature zone after saline **(B)**, capsaicin **(C)**, saline + capsaicin **(F)**, or gabapentin + capsaicin **(G)** administration, measured by EthoVision. Error bars represent SEM. **(D, H)** Comparison of mean preferred temperature between treatment groups showed capsaicin-treated mice preferred a cooler temperature than did saline-treated mice **(D)**. Conversely, gabapentin administration before capsaicin prevented heat aversion from occurring **(H)**. **(E, I)** Paired *t-*tests of individual mice’s mean preferred temperatures further reflected avoidance of higher temperatures in mice that received capsaicin **(E)**, whereas this was prevented in mice that received gabapentin **(I)**. Sal, saline; Cap, capsaicin; GBP, gabapentin. \**p* < 0.05, \*\*\**p* < 0.001

### Ferrule production and implantation for optogenetic light delivery

The surgical strategy used was adapted from the protocols for *in vivo* optogenetics described recently.^25, 27^ The prepared ferrule was sterilized with 70% ethanol and consisted of a 1.25□mm fiber optic ferrule (Ceramic ferrule, 1.2□mm diameter, 440□μm bore; Thor Labs, Germany) with a 400□μm fiber optic core (Multimode fiber, 400□≥ μm core, 0.39 NA; Thor Labs, Germany) that protruded out at a maximum length of 0.5 mm from the concave side of the ferrule. Mice were initially anesthetized using 5% isoflurane and then maintained on 2-2.5% isoflurane in a stereotactic frame. The back of the mouse was shaved and a 1.5 cm incision was made centered around the peak of the dorsal spinal curvature. Small spring scissors were used to make shallow incisions lateral to the T13 vertebrae which was identified based on its location relative to the peak of the dorsal spinal curve and the lowest rib. The vertebra was clamped using 2 pairs of transverse spinal adapters (Stoelting Co., Wood Dale, IL, USA) and the flesh was removed using spring scissors and fine forceps. Bleeding was controlled using autoclaved Kimwipes and absorbable gelatin sponges (Gelfoam, Pfizer Inc., New York, NY, USA). After ensuring stability of the vertebrae, a microdrill was used to flatten down the spinous process and slightly abrade the surface of the bone to promote adhesive bonding. The microdrill was then used to drill a hole in the center of the right side of the vertebra. Once spinal tissue was visible through the burr hole, the surgical site was cleared of any bone fragments and the prepared ferrule was lowered in place using stereotactic arms and cemented in place using Superglue and C&B Metabond (Parkell Inc., Brentwood, NY, USA). The skin around the implant was closed using a surgical adhesive (GLUture, Zoetis Inc., Parsippany-Troy Hills, NJ, USA). In the sham surgery animals, all surgical manipulation was performed except for the placement of the implant and its accompanying adhesives. The mice were allowed to recover for a minimum of 2 weeks prior to any behavioral testing.

### Optogenetic activation

The implant in the mouse was coupled to a patch cable (400ūμm core, 0.39 NA; Thor Labs, Germany) using a mating sleeve. The intensity and frequency of the 470 nm laser was controlled using a LED driver (Thor Labs) and a pulse width modulator (XY-LPWM, Protosupplies, USA), respectively. The laser output intensities reported here refer to the light intensity output from a test implant coupled to a patch cable through a mating sleeve. The output light intensity, measured using a PM100 light meter (Thor Labs), was calibrated with the current input from the LED driver. To determine the threshold light intensity for behavioral response, the patch cable was coupled to the implant while the mouse was under light isoflurane anesthesia. Each mouse was given 20 minutes to recover from the anesthetic and habituate to the fiber. The light intensity was increased in incremental steps of 0.2 mW with each intensity being presented at a maximum of 20 seconds or until a nocifensive response was observed. The first intensity level at which nocifensive responses like biting of the hind leg, fleeing and vocalization were first observed was classified as the threshold light intensity. Changes in thermal preference were assessed during acute peri-threshold stimulation and post suprathreshold stimulation of the TrpV1+ neurons. Acute peri-threshold stimulation consisted of photo-activation at a frequency of 10 Hz with intensity being set at 80% of the threshold light intensity. Suprathreshold stimulation was performed at 2 Hz at 200% of the threshold light intensity. 2 Hz light stimulation has been shown to produce optogenetically induced LTP in *ex vivo* spinal cord slices.^28^

### Behavioral Experiments

#### Rotarod

Evoked motor activity was tested in ferrule implanted and non-implanted sham surgery C57BL/6 mice using a rotarod (Bioseb, Chaville, France). Rotation was started immediately after the mice were placed on the rotarod, and accelerated from 4 to 40□RPM over a period of 120□s. The latency to fall was measured and a cut-off time of 120□s was set. Trials were repeated five times per mouse separated by 30□min each.

#### Open field

Spontaneous locomotion and anxiety related behaviors were assessed in ferrule implanted and non-implanted sham surgery C57BL/6 mice using an open field assay. The open field assay was conducted in a dimly illuminated square box. The floor of the box measured 30 × 30 cm. The mice were allowed to explore the novel box for 15 minutes while they were video recorded using an overhead video camera. The videos were later analyzed using EthoVision XT (Noldus Inc., Wageningen, Netherlands). The center point of the mouse was tracked, and total distance travelled, and time spent in the center zone of the assay was analyzed.

#### Mechanical sensitivity

Mechanosensitivity was assessed using von Frey filaments. Mice were placed in a small chamber with a grid floor and allowed to habituate for 30 minutes. Both hind paws were tested using the SUDO method^29^ and the paw withdrawal threshold was reported in pressure (g/mm^2^).^30^ This distinction in paw was relevant since the implant was always fixed on the right side of the vertebrae and we therefore expected possible lateralization in terms of behavioral or sensory deficit produced as the result of the implant.

### Data analysis

#### EthoVision parameters

EthoVision was used to calculate the cumulative amount of time each mouse spent in different temperatures across the arena. The 36.0 cm length of the arena was separated into eighteen 2.0 cm temperature zones (**Fig. 1C**). Heatmaps were generated along with coordinates of each mouse’s centre-point position, hereafter referred to as centre, within the arena (**Fig. 1D**). Coordinate samples were taken once per second for each 30-minute trial, for a total of 1800 coordinates per trial. Raw coordinates were given in centimetres and converted into pixels for comparison with DeepLabCut’s raw coordinates.

#### DeepLabCut parameters

DeepLabCut (version 2.0.4.1, http://www.mousemotorlab.org/deeplabcut) was used to validate EthoVision’s tracking data (**Fig. 1D**). DeepLabCut is an open-source animal tracking program that uses deep neural networks to recognize labelled points in videos.^16^ DeepLabCut’s tracking algorithm was trained to recognize the mouse’s centre, labelled as visually close to EthoVision’s calculated centre as possible, with 200 distinct marked reference frames. All other settings were left as default as specified in the setup protocol.^16, 31^ No further refinement of tracked points were performed after training. All training and video analyses were done using an NVIDIA GeForce RTX 2080Ti supported by an Intel® Core^TM^ i7-4790 and 16 GB of RAM, on Ubuntu 18.04.2. Similarly, 1 coordinate sample was taken per second for each 30-minute trial by averaging the positions for all 30 frames each second, for a total of 1800 coordinates per trial. Raw coordinates were obtained in pixels and converted into centimetres and zones for further validation of EthoVision’s raw coordinates. All trial recordings were recorded at 30 frames per second and normalized to a 16:9 aspect ratio, 1280×720 resolution using a scaling factor for data analysis.

#### Statistical analysis

Mean temperature preferences were determined by fitting data to a Gaussian distribution with least squares regression and no special handling of outliers. These means of the controls and treatments were then compared through a parametric paired two-tailed *t*-test. Any unpaired mice were excluded from the calculation of statistical significance. The performance of implanted and sham surgery animals in behavioral experiments was compared using unpaired two-tailed *t*-tests and repeated measures two-way analysis of variance (ANOVA). Paired two-tailed *t*-tests and repeated measures one-way ANOVA were used to compare means in the behavioral experiments conducted to validate our optogenetic stimulation protocols. All statistical analyses were performed in GraphPad Prism 8, with a significance level set at P < 0.05. Data are presented as mean ± standard error of the mean (SEM).

## Results

### Arena calibration

To confirm the arena had a linear temperature gradient, the temperature of the arena floor was measured every 2.0 cm along its length with an infrared thermometer every 5 minutes for 1 hour. Once temperatures of the floor stabilized, the cold, middle, and hot ends measured temperatures of 11.0 ± 0.6 °C, 29.0 ± 0.9 °C, and 51.0 ± 0.7 °C, respectively, and a clear linear gradient was observed along the length of the arena (**Fig. 1B**).

### Capsaicin shifted temperature preference towards lower temperatures

We first used the thermal preference assay to examine whether intraplantar capsaicin would produce a change in mouse thermal preference. We first performed an intraplantar injection of either saline or capsaicin into the mouse hind paw 30 minutes prior to testing in a randomized crossover manner (**Fig. 2A**). We found that mice injected with saline showed a preference for a temperature of 32.7 ± 0.3 °C (**Fig. 2B**), whereas those treated with capsaicin preferred a slightly cooler temperature of 31.3 ± 0.3 °C (**Fig. 2C**). Comparing both groups via Gaussian fit of each treatment group’s summed positional data over the experiment revealed a small but significant difference between preferred temperatures across the two treatment groups, with capsaicin-treated mice preferring cooler temperatures (**Fig. 2D**; *F*_(3,264)_ = 8.530, *p* < 0.0001). Moreover, this shift to cooler temperatures was seen within mice with paired analysis of mean preferred temperature after each treatment, indicating that mice consistently showed a preference for cooler temperatures after capsaicin was administered (**Fig. 2E**; Paired *t*-test, *t*_5_ = 3.521, *p* = 0.017).

### Gabapentin prevented capsaicin-induced heat aversion

To assess if the capsaicin-induced avoidance of higher temperatures was a result of pain associated with capsaicin injection rather than a systemic change in thermal preference we administered either saline or a non-opioid analgesic, gabapentin, 30 minutes prior to capsaicin injection (**Fig. 2A**). Mice that received capsaicin preferred 31.2 ± 0.4 °C (**Fig. 2F**), whereas mice that received gabapentin and capsaicin preferred 32.4 ± 0.3 °C (**Fig. 2G**). Comparing these two groups with a Gaussian fit of the summed treatment group positional data similarly revealed a small but significant difference between their mean preferred temperatures, with gabapentin and capsaicin treated mice preferring higher temperatures compared to mice that were only treated with capsaicin (**Fig. 2H**; *F*_(3, 264)_ = 3.745, *p* = 0.012). As before, this difference in temperature preference was observed within mice as a paired analysis consistently revealed a preference for higher temperatures after treatment with gabapentin (**Fig. 2I**; Paired *t*-test, *t*_5_ = 3.063, *p* = 0.028).

### DeepLabCut provides an open-source approach to the assessment of mouse thermal preference

The positional analysis completed thus far employed EthoVision, which is a powerful and commonly used commercial behavioral analysis program. To further lower the cost of our system, we investigated whether an open source tracking software, DeepLabCut, could be used to detect positional differences described in the thermal preference assays above in place of EthoVision.

We first confirmed that DeepLabCut could track the position of the centre of the mouse similarly to EthoVision. The DeepLabCut training labels were created as visually close to EthoVision’s centre as possible before training its tracking algorithm. After training the network with >1,000,000 iterations, the network was able to detect the center of the mouse with an accuracy of ± 4.32 pixels (± 0.17 cm) in test videos compared to EthoVision. The mean temperature preference for each mouse tested above was then determined by EthoVision and DeepLabCut. Across all these mice there was an extremely high correlation between the mean preference assessed with the two methods (**Fig. 3A**; Pearson’s *r* = 0.94), indicating that DeepLabCut can be used to assess mouse thermal preference. Moreover, we detected no significant differences in the mean temperature preference of any treatment group when comparing between analysis approaches (**Fig. 3B, C**)

**Figure 3.**
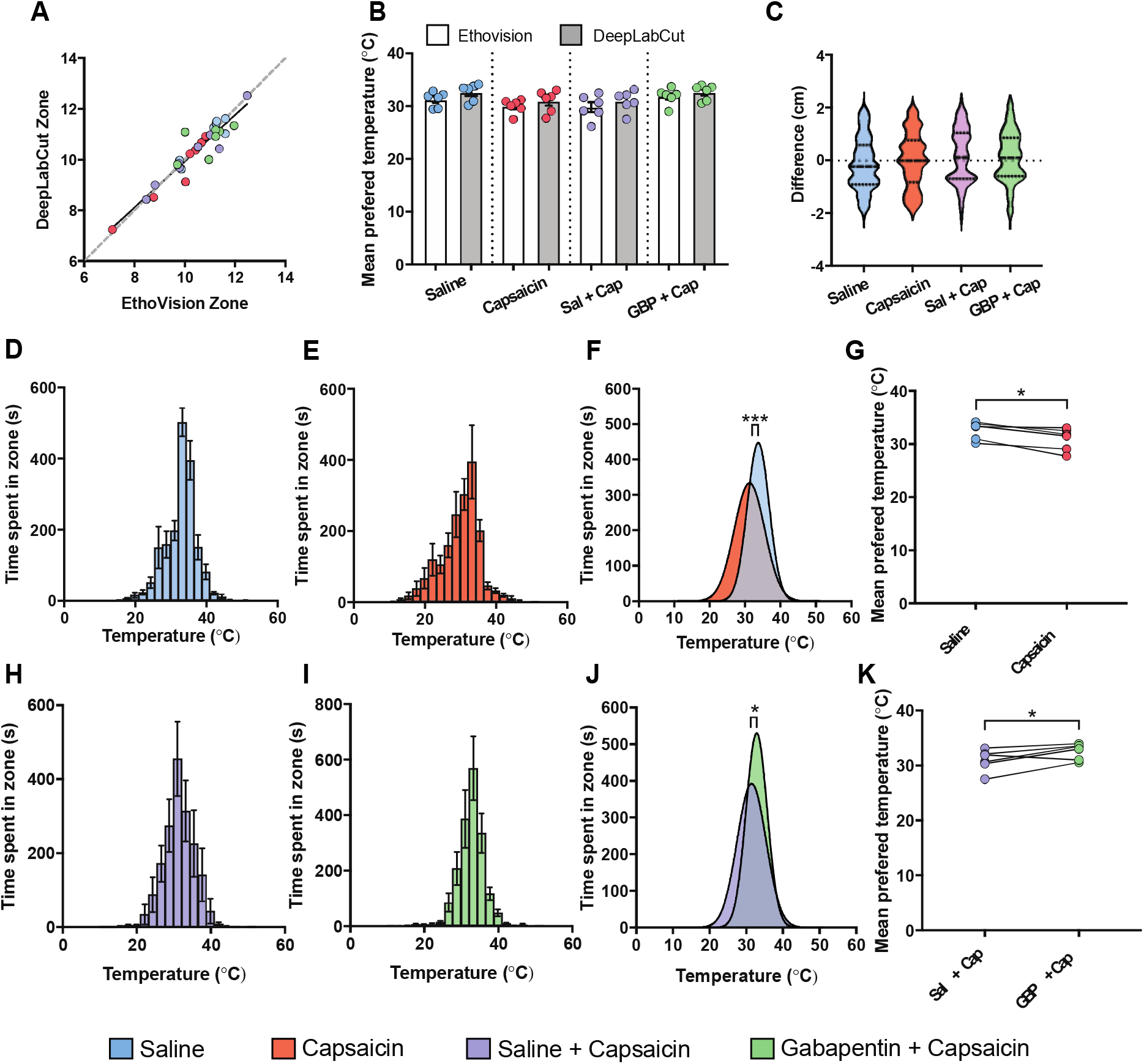
DeepLabCut and EthoVision provided equal animal tracking precision. **(A)** Comparison of each mouse’s mean preferred temperatures (*n* = 31) determined by DeepLabCut and EthoVision showed that DeepLabCut could track mice’s centres with high precision, comparable to that of EthoVision (Pearson’s *r* = 0.94). **(B, C)** There was no difference in mean preferred temperature obtained for each of the groups by both software using a one-way ANOVA. **(D, E, H, I**; *n* = 6) Total time spent in each temperature zone after saline **(D)**, capsaicin **(E)**, saline + capsaicin **(H)**, and gabapentin + capsaicin **(I)** administration, measured by DeepLabCut. Error bars represent SEM. **(F, J)** Comparing mean preferred temperatures measured by DeepLabCut validated that capsaicin indeed shifted temperature preference to cooler temperatures **(F)** and that gabapentin prevented this from occurring **(J). (G, K)** Paired *t-*tests of individual mean preferred temperatures reflected a shift in preference to cooler temperatures post-capsaicin administration **(G)** that was no longer seen if gabapentin was given before capsaicin **(K)**. Sal, saline; Cap, capsaicin; GBP, gabapentin; ANOVA, analysis of variance. \**p* < 0.05, \*\*\**p* < 0.001

To compare approaches at a pixel-level of tracking in all samples, we then calculated the distance between EthoVision and DeepLabCut’s predicted centre location for every frame in the first trial of each treatment group (**Fig. 3C**). We found that the 95% confidence interval of the median difference was −0.048 cm to 0.021 cm, with a minimum and maximum difference in centre points of −2.666 and 2.620 cm, respectively. Negative values represent samples where DeepLabCut’s predictions visually fell to the left of the mouse, while positive values are to the right. Since 0 fell within the confidence interval and even the most extreme differences did not exceed the visual size of the mice, these data indicate that DeepLabCut largely provided the same frame-by-frame precision observed with EthoVision.

Finally, we repeated the positional analysis of mice treated with saline or capsaicin, and capsaicin or capsaicin + gabapentin (**Fig. 2**). As seen with EthoVision, our positional analysis with DeepLabCut revealed that mice treated with capsaicin preferred cooler temperatures than mice treated with saline (**Fig. 3D-G**), and that mice treated with gabapentin and capsaicin preferred warmer temperatures than those treated with saline and capsaicin (**Fig. 3H-K**). These data further support that DeepLabCut provides an open-source option for the analysis of mouse thermal preference as assessed using the low-cost thermal preference assay described here.

### Optogenetic activation of the central terminals of TrpV1+ afferents did not recapitulate changes in thermal preference induced by intraplantar capsaicin

Having established that capsaicin can produce an aversion to warmer temperatures we sought to assess whether optogenetic activation of the central terminals of the TrpV1+ afferents would similarly shift thermal preference. To deliver light to our region of interest the ferrule implant was fixed in the T13 vertebra that overlies a portion of the L4-L6 spinal cord region that receives innervations from the hindlegs of the mouse (**Fig. 4A**).The implant was well tolerated by mice, with most animals retaining implants for a minimum of 2 months. To ensure that the implant had limited impact on mouse motor and sensory function, we assessed evoked motor activity, spontaneous locomotion and mechanical sensitivity using rotarod, open field and von Frey filaments, respectively, in ferrule implanted and sham operated C57BL/6 mice. There was no difference in performance on the rotarod between sham and implanted animals (**Fig. 4B**; Repeated measures two-way ANOVA, effect of treatment: *F*_(1, 4)_ = 0.149, *p* = 0.72, effect of trial: *F*_(1.8, 7.2)_ = 4.078, *p* = 0.070, effect of interaction: *F*_(4, 16)_ = 0.516, *p* = 0.73). There was also no difference in distance travelled in the open field assay (**Fig. 4C**; Unpaired *t*-test, *p* = 0.32). Overall these data indicate there was no difference in both evoked and spontaneous motor activity between the two experimental groups. Additionally, time spent in the center zone was analyzed in the open field to assess for thigmotaxis, an anxiety related behavior that can confound results of a preference-based assay like the one described within this paper. No differences were found for time spent in the center zone of the open field between implanted and sham mice (**Fig. 4D**; Unpaired *t*-test, *p* = 0.62). Finally, no differences were found in the paw withdrawal thresholds of the ipsilateral and contralateral paws between the surgery and the sham animals, indicating that the implant did not change nociceptive thresholds or cause sensory deficit (**Fig. 4E**; Repeated measures two-way ANOVA, effect of treatment: *F*_(1, 4)_ = 0.166, *p* = 0.71, effect of paw: *F*_(1, 4)_ = 0.339, *p* = 0.59, effect of interaction: *F*_(1, 4)_ = 0.003, *p* = 0.96). We used implanted TrpV1-ChR2 mice for the following experiments.

**Figure 4.**
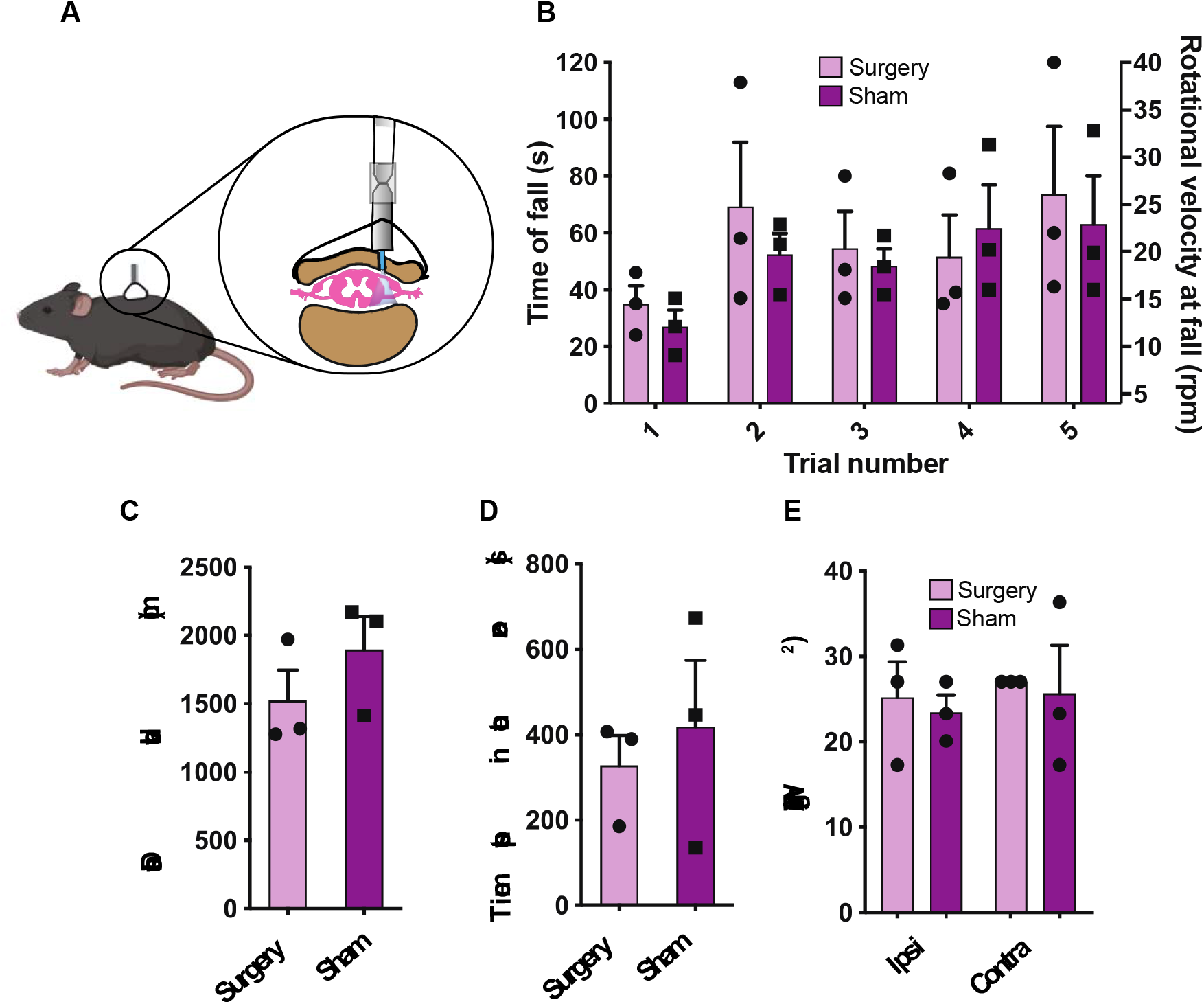
The ferrule implant surgery did not impact mouse sensory or motor behavior. **(A)** A cartoon illustration of the surgical strategy employed which depicts the placement of the implant relative to the vertebrae and spinal cord. **(B)** Evoked motor performance did not differ between implant surgery and sham surgery animals and was compared using a repeated measures two-way ANOVA. **(C)** No difference in distance travelled by implant surgery and sham surgery animals in open field assay as compared using an unpaired t-test. **(D)** No difference in time spent in center zone by implant surgery and sham surgery animals in open field assay as compared using an unpaired t-test. **(E)** Based on a repeated measures two-way ANOVA PWT of the ipsilateral and contralateral paws of the implant surgery and sham surgery animals did not differ. *n* = 3 per group. Error bars represent SEM. ANOVA, analysis of variance; PWT, paw withdrawal threshold.

To assess whether stimulation of TrpV1+ fibers is sufficient to induce changes in thermal preference, mice were connected to the LED light source and placed on the preference assay. Two different stimulation paradigms were used: (1) to assess whether peri-threshold stimulation can produce acute changes in thermal preference mice were initially placed on the assay for 15 minutes with the LED off to establish baseline preference, after which perithreshold stimulation (10Hz, 80% threshold intensity) was then initiated for another 15 minutes (**Fig. 5A**); (2) to assess whether stimulation at suprathreshold light intensities to initiate central sensitization would change temperature preference. In this paradigm, mice were placed on the assay for 15 minutes, after which they were removed from the assay and placed in an altered context where they received suprathreshold stimulation (2 Hz, 200% threshold) for 10 minutes followed by a 30-minute wait time within that same context in accordance with previous studies of optogenetic sensitization.^26, 28^ The mice were then placed back on the assay for 15 minutes for assessment of temperature preference (**Fig. 5B**).

**Figure 5.**
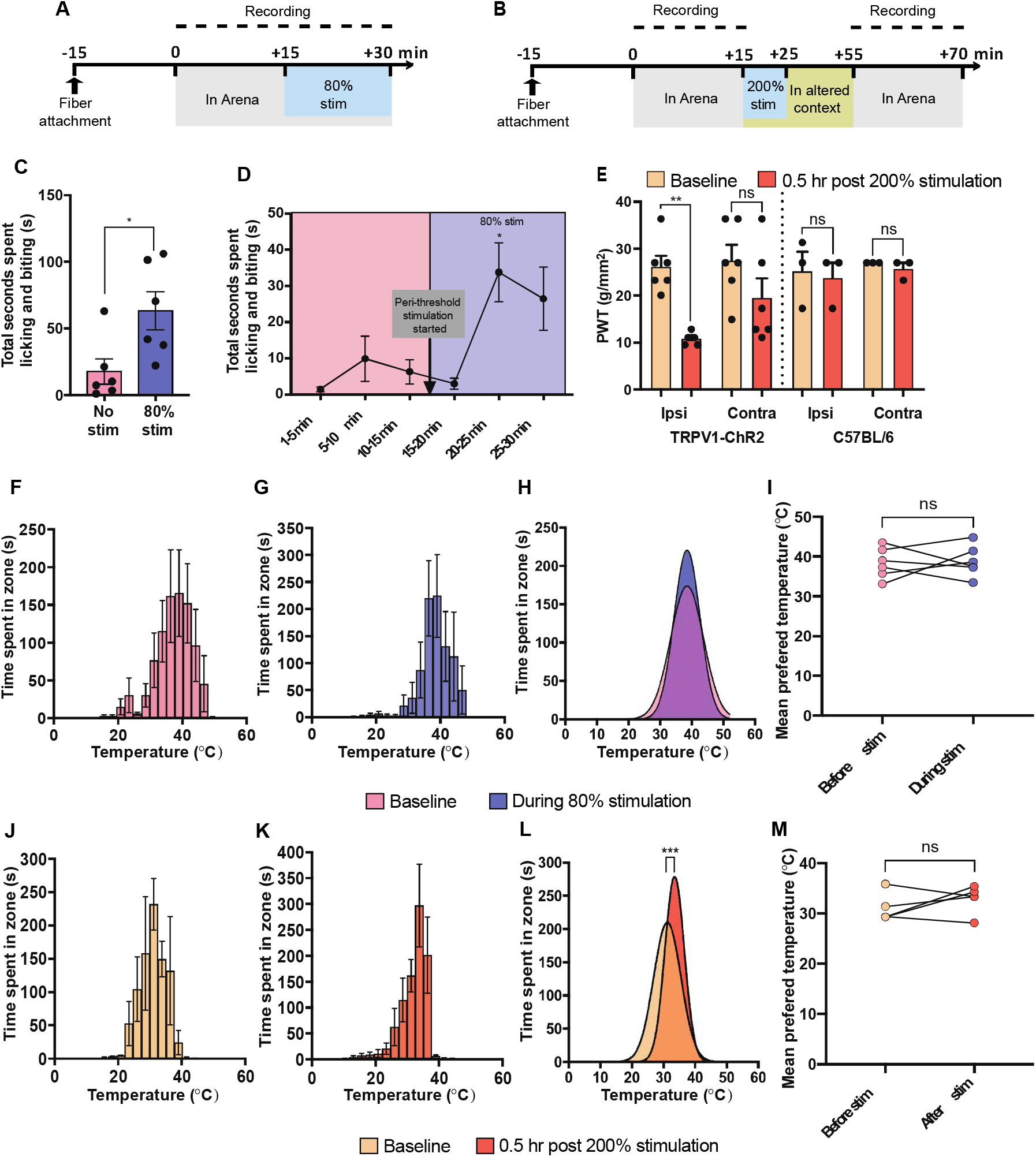
Optogenetic activation of the central terminals of the TrpV1+ afferents did not produce a shift in temperature preference. **(A)** Experimental timeline for assessing temperature preference before and during peri-threshold activation of TrpV1+ neurons. **(B)** Experimental timeline for assessing temperature before and after suprathreshold activation of TrpV1+ neurons. **(C)** Mice spent more time licking and biting their hindquarter during peri-threshold stimulation than under baseline and (D) these nocifensive behaviors emerged after more than 5 minutes of the peri-threshold stimulation being active (*n* = 6). **(E)** Suprathreshold stimulation caused a decrease in the paw withdrawal threshold of the ipsilateral paw in only the TrpV1-ChR2 mice (*n* = 3-6 per group). **(F, G)** Cumulative time spent in each temperature zone during **(F)** Pre-stimulation and **(G)** during stimulation for the perithreshold stimulation experiments calculated using Ethovision. **(J, K)** Cumulative time spent in each temperature zone during **(J)** Pre-stimulation and **(K)** after stimulation for the suprathreshold stimulation experiments calculated using Ethovision. **(H)** Comparison of the peaks of the Gaussian regressions fit to the aggregated data from all mice during pre-stimulation and stimulation conditions failed to reveal a shift in temperature preference during perithreshold stimulation. **(I)** A paired t-test of temperature preferences of the individual mice, obtained through gaussian fits to individual mouse data, also did not show a difference in preference under peri-threshold stimulation (*n* = 6). **(L)** Comparison of the peaks of the Gaussian regressions fit to the aggregated data from all mice during prestimulation and post-stimulation conditions revealed that mice on average preferred a warmer temperature post suprathreshold stimulation. **(M)** However, a paired t-test of mouse specific temperature preferences did not uphold this trend and no differences were found in temperature preference pre- and post-suprathreshold stimulation (*n* = 5). \**p* < 0.05, \*\**p* < 0.005, \*\*\**p* < 0.001; ns, not significant.

The video recordings of the mice on the assay for the peri-threshold thermal assay experiments were manually scored for nocifensive licking and biting behavior. We found that mice spent more time engaging in licking and biting behavior during peri-threshold stimulation compared to the same mice prior to stimulation (**Fig. 5C**; Paired *t*-test, *p* = 0.02). Notably, the increased nocifensive behavior only emerged after the five minutes of perithreshold stimulation (**Fig. 5D**; Repeated measures one-way ANOVA, *F*_(2.6, 13.1)_ = 7.29, *p* = 0.01). For the TrpV1-ChR2 mice stimulated with peri-threshold stimulation, mice had a thermal preference of 38.3 ± 0.7 °C at baseline (**Fig. 5F**) prior to stimulation and a preference of 38.6 ± 0.6 °C during peri-threshold optogenetic stimulation of TrpV1+ afferents (**Fig. 5G**). Comparing the summed positional data of the animals at baseline and during perithreshold stimulation revealed no difference in the gaussian curves fit of the data sets (**Fig. 5H**; *F* _1,210)_ = 0.102, *p* = 0.75). Similarly, no within mouse change in thermal preference was observed before and during peri-threshold stimulation (**Fig. 5I**; *t*_5_ = 0.209, *p* = 0.84), indicating that acute excitation of TrpV1+ afferents does not modify thermal preference.

In order to confirm suprathreshold stimulation induced sensitization, we assessed mechanical paw withdrawal thresholds using von Frey filaments in TrpV1-ChR2 and C57BL/6 mice that received suprathreshold stimulation. The suprathreshold stimulation intensity used for C57BL/6 mice the average suprathreshold threshold of all TrpV1-ChR2 mice. Suprathreshold stimulation caused a decrease in the paw withdrawal threshold of the ipsilateral paw of the TrpV1-Chr2 mice but not the C57BL/6 mice (**Fig. 5E**; Repeated measures one-way ANOVA, *F*_(1.9, 9.8)_ = 6.878, *p* = 0.01) indicating that the change in mechanical sensitivity was not a non-specific effect of blue light. Having established that suprathreshold stimulation successfully induced mechanical hypersensitivity we sought to assess whether it also induced changes in thermal preference. Overall, TrpV1-ChR2 mice exhibited a mean thermal preference of 31.2 ± 0.5 °C at baseline (**Fig. 5J**) and 33.5 ± 0.3 °C after suprathreshold stimulation (**Fig. 5K**). Comparing the summed positional data of the animals at baseline and after suprathreshold stimulation revealed that sensitization by optogenetic activation of central terminals of TrpV1+ afferents unexpectedly produced a change in the overall positional distribution of animals towards warmer temperatures (**Fig. 5L**; *F*_(1,174)_ = 12.490, *p* < 0.001). This was in contrast to the opposite trend observed as a part of the capsaicin experiments. However, a paired t-test of the individual mouse temperature preferences failed to hold the same trend as the cumulative gaussian fits (**Fig. 5M**; *t*_4_ = 1.118, *p* = 0.33).

## Discussion

Our findings show that an automated thermal selection assay can reveal subtle changes in thermosensory phenotypes in mice. We detected small, yet robust, differences in temperature preference pre- and post-capsaicin treatment across very short distances in a minimally invasive manner and without extensive investigator intervention. Our results also show that the behavioral phenotype, with respect to sensory hypersensitivity, differs in mice injected with capsaicin, a TrpV1 agonist, versus those that have TrpV1+ neurons activated optogenetically. We also validated DeepLabCut as a precise animal tracking program and, for the first time, demonstrated its applicability in pain research. Overall, the simplicity of our assay and ease of access to the materials and technologies used support its use as an easily implemented assay for investigation of new analgesic drugs.

The significant translational gap in pain research has promoted increased characterization of non-reflexive indicators of spontaneous pain, as these may be of greater translational value.^32^ As opposed to stimulus-evoked pain, spontaneous pain does not require external stimuli triggers and is seen in almost all chronic pain conditions.^33^ Unlike nocifensive reflexes that can be evoked within the spinal cord or subcortical circuits^7, 10, 11^; chronic pain involves a distributed nociceptive network that integrates supraspinal regions that control affective, cognitive, memory, and motor functions along with spinal cord nociceptive processing.^34^ Therefore, a comprehensive preclinical assessment of pain would include measures for both acute and spontaneous behavioral responses to noxious and innocuous stimuli. Recent studies have characterized a variety of mouse behaviors that may be more reflective of spontaneous pain and supraspinal processes associated with clinical conditions, including free-choice temperature preference assays.^6, 7, 9, 12–15^ However, most assays used to assess thermal preference rely on mouse avoidance responses to noxious temperatures rather than assessment of native preference.^7^ In contrast, our assay does not necessarily expose mice to noxious temperature ranges and therefore provides a measure of response to innocuous temperature, allowing for a study of thermal allodynia or thermotaxis.

The selection of a preferred temperature by mice likely involves a complex evaluative process that integrates body temperature, environmental temperature, and the modulation of temperature sensation by pathological conditions, such as peripheral or central sensitization.^35, 36^ Thus, in direct contrast to reflexive responses in acute nociception assays, the preference of a particular temperature is driven by persistent sensory processing and the motivation to move to different temperatures may reflect non-reflexive and integrative sensory processing or coping behavior with important motivational or homeostatic components, such as avoiding pain or discomfort.^37, 38^ Additionally, given that mice are a prey species that tend to hide overt behavioral signs of pain or distress^12,15^, assays that rely on more subtle behavioral outputs, such as thermal selection, may provide a unique insight into subtle but physiologically relevant changes in sensory preferences and thresholds.

The intraplantar injection of capsaicin is associated with thermal and mechanical sensitization.^17–19^ Our findings that intraplantar capsaicin injection induces a preference for cooler temperatures are consistent with these findings, as thermal sensitization and thermal allodynia would be expected to produce an aversion to warm temperatures. Activation of TrpV1 receptor is not necessary for the expression of thermal preference as previous research has shown that TrpV1 knockout mice display normal behavior in thermal gradient assays.^36, 39^ This is not unexpected as TrpV1 is activated by noxious temperatures and we mice did not observe mice actively exploring regions of the assay at or above noxious temperature thresholds in this study.

However, while TrpV1 knockouts are mostly indistinguishable from their wild type counterparts in innocuous thermal sensitivity they fail to develop thermal allodynia and hyperalgesia during inflammation^40, 41^, implying an increased role for TrpV1 activity in inflammation. Indeed, TrpV1 has a lower thermal threshold in cases of inflammatory pain and this effect is likely mediated by phosphorylation-dependent upregulation of channel function through inflammatory mediators.^42–44^ TrpV1 has multiple potential targets sites for phosphorylation by proteins kinases. There is a large body of evidence to support the role of protein kinase C (PKC) and protein kinase A (PKA) signalling pathways in the sensitization of TrpV1.^42–44^ PKC, activated via bradykinin, phosphorylates TrpV1 and potentiates heat-evoked responses by lowering the threshold temperature for channel activation.^43, 45–47^ TrpV1 is also known to be sensitized through the prostaglandin mediated activation of PKA.^48, 49^ Capsaicin has been shown to result in peripheral neurogenic inflammation.^50^ This is often characterized by the sustained release of an “inflammatory soup” which most notably consists of the peptides Substance-P and C Calcitonin gene-related peptide (CGRP), that are released from peripheral terminals of primary afferents that trigger the release of additional mediators from non-neuronal cells such as bradykinin, prostaglandins and neurotrophins (NGF).^51^ Overall, the sensitization of TrpV1 renders TrpV1+ afferents more excitable and lowers their thermal threshold of activation. This peripheral inflammatory process likely contributed to the change in thermal preference of mice after intraplantar injection of capsaicin. In support of this, we observed that gabapentin reduced thermal preference of mice injected with capsaicin. Gabapentin exerts part of its analgesic properties through its ability to reduce the presence of inflammatory mediators.^52^ As a result, this mechanism may explain how gabapentin was able to reverse the effect of capsaicin-induced warm-temperature aversion.

We unexpectedly observed that acute peri-threshold optogenetic activation of TrpV1+ afferent terminals in the spinal cord did not alter thermal preference. This finding suggests that the central activation of TrpV1+ afferents itself is insufficient to induce changes in thermal preference, and that this behavioral outcome requires a host of other changes beyond an increase in the excitability of TrpV1+ afferents. This summary of findings above indicates that peripheral inflammation induced by capsaicin is a crucial process by which thermal sensitization arises. Although not quantified, we also observed a lack of visual signs of inflammation such as redness and edema in the hind paws of the stimulated TrpV1-ChR2 implanted animals, as is typically seen in response to capsaicin. Notably, previously studies optogenetically activating hindleg nociceptors have also failed to produce neurogenic inflammation.^23,28^ Specifically, channelrhodopsin mediated activation of Na_v_1.8^28^ and TrpV1^23^ primary afferents has been shown to not produce neurogenic inflammation. We speculate that the likely failure to produce peripheral neurogenic inflammation in this study may further arise from the fact that we were activating central projections of the primary afferent rather than peripheral terminals.

The administration of intraplantar capsaicin also causes mechanical hyperalgesia arising from central sensitization.^53^ It is possible that central sensitization further contributed to the change in thermal preference observed after capsaicin injection. However, we did not observe a change in thermal preference towards lower temperatures after induction of sensitization by suprathreshold activation of central TrpV1-ChR2 terminals, further supporting the argument the changes in thermal preference caused by intraplantar capsaicin injection arises from peripheral mechanisms. Alternatively, it is possible that the biophysical differences between TrpV1 and ChR2 channels underlie the inconsistencies in behavioral responses to capsaicin and optogenetic afferent activation: TrpV1 is a non-selective, large pore cation channel with a notable calcium permeability^54^, while ChR2 is a lower conductance cation channel with lower calcium permeablity.^55^ This raises the possibility that the lack of calcium influx in the optogenetic stimulation paradigm is a factor in the lack of behavioral changes. Additionally, optogenetic activation of afferent terminals may not fully recapitulate the pattern neuronal activity induced by capsaicin. However, the induction of mechanical sensitization we observed in response to central TrpV1-ChR2 activation implies that our optogenetic protocol is capable of recapitulating the central sensitization associated with intraplantar capsaicin injection.

Usage of automation in pain research is steadily increasing. Grimace Scales for mice and rats have now been paired with deep neural networks and specialized software, respectively, for recognizing subtle facial expressions that reflect spontaneous pain with accuracy comparable to a trained human researcher.^56, 57^ In our case, DeepLabCut is a new open-source application that uses pre-trained deep neural networks to track digitally inserted points in video recordings. The program’s deep-learning algorithm allows researchers to quickly train the program to recognize novel subjects in highly variable environments without the need of physical markers on the subjects themselves.^16, 31^ Here, we showed that DeepLabCut could track mice comparably to EthoVision using only 50 more reference frames to train the network than the default 150. Even without further refinement after the first iteration of training, there was high fidelity and consistency in tracking our desired points upon visual inspection on a frame-by-frame basis. However, as the program is limited to only tracking coordinates of individual points, more intricate software or algorithms must be developed to characterize complex subject behaviors or when working with multiple points. Nonetheless, DeepLabCut’s versatility, high generalizability to novel settings, and near-human accuracy in tracking will indeed make it a valuable asset for future automated research.

Our assay had a temperature range from approximately 10 °C to 50 °C so as to allow for possible exposure of the mice to noxious temperatures. As expected, mice did not explore zones in the assay that had aversively high or low temperatures but stayed within a preferred range of temperatures consistent for mice. Reducing the total temperature range of the assay would possibly allow for more nuanced or precise analysis of temperature preference. Notably, different strains, sexes, ages, and housing conditions of mice can affect thermal preference.^37, 58, 59^ For example, older (>11 months) CD-1s prefer warmer temperatures, whereas older C57BL/6 mice do not. Singly housed older CD-1s also prefer warmer conditions than group housed ones, while younger CD-1s did not differ in preference when housed individually or in groups. Furthermore, female mice also tend to prefer warmer temperatures than males, likely attributed to differing metabolic needs, body composition, and hormonal physiology. Our assay would allow further research of such physiological variables in temperature preference in addition to the pathological variables examined here. Given that strain differences in thermosensory and thermoregulatory processes^38, 60^ have been reported, we acknowledge that mouse strain may be a confounding factor for the difference in phenotypic response observed in response to capsaicin injections in CD-1 mice and optogenetic activation of TrpV1+ neurons in TrpV1-ChR2 C57BL/6 mice.

Overall, we report the development and validation of an easily sourced assay for the automated assessment of thermal preference in mice. Using this assay, we confirmed changes in thermal preference induced by peripheral capsaicin injection, and further contrasted these findings against the lack of a change in thermal preference induced by optogenetic stimulation of TrpV1+ afferents at the level of the spinal cord. These findings support the use of this assay to investigate central and peripheral mechanisms governing thermal preference in freely behaving animals.

## Acknowledgements

This research was supported by funding from the Natural Sciences and Engineering Research Council of Canada (RGPIN-2016-05538 to RPB), the Canadian Institutes of Health Research (FRN 162179; RPB), a Canada Research Chair in Sensory Plasticity and Reconsolidation.the Canadian Pain Society (RPB), and the University of Toronto Centre for the Study of Pain (MZ, RPB).

## Author Contributions

Pharmacological experiments and DeepLabCut animal tracking were performed by JL, while all optogenetic experiments and associated animal tracking were performed by MZ. All authors planned the experiments, wrote, and edited the final manuscript.

